# Computational Methods for Optimal Coil Placement and Maximization of Lobule-Focused Cortical Activation in Cerebellar TMS

**DOI:** 10.1101/2025.02.27.640697

**Authors:** Xu Zhang, Roeland Hancock, Sabato Santaniello

**Author notes:** **<u>Correspondence:</u>** Sabato Santaniello, 260 Glenbrook Road, Unit 3247, University of Connecticut, Storrs, CT 06269-3247 (USA). **AUTHOR CONTRIBUTIONS:** X.Z.: Conceptualization, Data Curation, Formal Analysis, Investigation, Methodology, Software, Validation, Visualization, Writing (Original Draft Preparation). R.H.: Conceptualization, Methodology, Resources, Supervision, Writing (Review & Editing). S.S.: Conceptualization, Funding Acquisition, Methodology, Project Administration, Resources, Supervision, Visualization, Writing (Review & Editing).

## Abstract

**Background:** Coil placement on the cerebellum lacks accuracy in targeting the intended lobules and limits the efficacy of cerebellar transcranial magnetic stimulation (TMS) in treating movement disorders.

**Objective:** Develop a multiscale computational pipeline and method to rapidly predict the cellular response to cerebellar TMS and optimize the coil placement accordingly for lobule-specific activation.

**Methods:** The pipeline integrates 3T T1/T2-weighted MRI scans of the human cerebellum, lobule parcellation, and finite element models of the TMS-induced electric (E-) fields for figure-of-eight coils (MagStim D70) and double-cone coils (Deymed 120BFV). A constrained optimization method is developed to estimate the fiber bundles from cerebellar cortices to deep nuclei and, for both coil types, find the coil placement and orientation that maximize the E-field intensity in a user-selected lobule. Multicompartmental Purkinje cell models with realistic axon geometries and Gaussian process regression are added to predict the recruitment in the Purkinje layer.

**Results:** Our pipeline was tested in five individuals to target the left lobule VIII and resulted in normalized E-field intensities at the target 49.6±25.6% (D70) and 29.3±17.7% (120BFV) higher compared to standard coil positions (i.e., 3 cm left, 1 cm below the inion), mean±S.D. The minimum pulse intensity to recruit Purkinje cells on a 4 mm^2^-surface in the target decreased by 21.6% (range: 4.7-55.0%) and 10.7% (range: 7.9-18.2%), and the spillover to adjacent lobules decreased by 70.6±16.3% and 71.7±20.8% compared to standard positions (D70 and 120BFV, respectively).

**Conclusion:** Our tools are effective at targeting specific lobules and pave the way toward patient-specific setups.

## INTRODUCTION

Transcranial magnetic stimulation (TMS) of the cerebellum is an effective tool to restore motor function in patients with ataxia [1, 2], improve dysphagia and swallowing impairments secondary to stroke [3, 4], reduce levodopa-induced dyskinesia in patients with Parkinson’s disease [5], and suppress essential tremor [6, 7]. Therapeutic outcomes, however, have been variable [8], with the effects and tolerability of cerebellar TMS being highly dependent on the coil configuration and target selection [9, 10].

Most clinical studies that target the motor cerebellum have used figure-of-eight planar coils placed either 3 cm lateral to the inion (*3L*) or, more commonly, 3 cm lateral and 1 cm inferior to the inion (*3L1I*), e.g., [1, 5–7, 11]. It is unclear, though, how much of the motor area of cerebellum is reached by placing the coil in position *3L* or *3L1I*. Direct measurable outputs of cerebellar TMS are lacking [12], and computational studies [13–16] have shown that the effects of TMS critically depend on the cerebellar anatomy, tissue composition, and distance from the surface, thus suggesting that the cerebellar motor area is hardly reached when coils are positioned in *3L* and *3L1I*. Also, planar coils hardly reach targets below the superficial tissues and have inferior performance in eliciting cerebellar-brain inhibition compared to double-cone coils [9, 10], thus suggesting that non-planar coils could be more advantageous in cerebellar TMS compared to planar coils. However, the best position and orientation of non-planar coils on the cerebellum are unknown, and a generalizable approach to coil placement optimization for non-planar coils in cerebellar TMS is needed to facilitate the development of more effective therapeutic applications.

Studies [17–19] have proposed to optimize the coil placement offline by calculating the TMS-induced electric field (E-field) via finite element methods (FEM) in a desired region of interest (ROI) for various combinations of coil position and orientation and then selecting the combination that maximizes the E-field intensity in the ROI. These methods, though, were mainly developed for TMS of the cerebral cortex and hardly translate to cerebellar TMS because of the cerebellum’s suboccipital location, larger distance from the scalp, and different neuronal arrangement compared to the cerebrum [9, 20–23]. Studies [24–27], instead, have used MRI-guided navigation and probabilistic mapping to minimize the coil-to-ROI distance online, thus maximizing the average E-field intensity within the ROI. The E-field intensity alone, however, is not necessarily predictive of the effects on neurons in any cerebellar ROI due to the complex local geometry of the gyri [13, 28], thus requiring additional methods to assess the cellular response to TMS. Finally, the cerebellar projection neurons, i.e., the Purkinje cells, have long axons that run toward the deep cerebellar nuclei and have lower activation threshold compared to soma [29–32], thus suggesting that TMS can induce action potentials well underneath the cerebellar cortex. This, however, poses further challenges to the selection of effective coil positions.

To address these issues, we developed a computational pipeline that estimates the cellular response to cerebellar TMS pulses both rapidly and accurately and, for any assigned cerebellar ROI (e.g., a segmented lobule), quickly calculates the best coil placement to enhance the cellular activation within the ROI. First, we propose a new machine learning algorithm to rapidly predict the activation thresholds of neuronal fibers stemming out of the Purkinje layer based on the TMS-induced E-fields. Differently from recent algorithms for neural fiber activation prediction [33–36], which require extensive training and numerical simulations, our solution needs limited training as it combines Gaussian process regression [37] and feature computation. Our features are designed to capture the spatial variation of TMS-induced E-fields along the primary axons of the Purkinje cells, and the axon development is obtained via constrained optimization on MRI-derived white matter pathways. Secondly, we develop a novel optimization algorithm to rapidly determine the best coil position and orientation constrained to the cerebellar anatomy that maximizes the E-field intensity in the ROI. Finally, we combine these algorithms with a multiscale framework that calculates the response of multicompartmental Purkinje cell models to TMS-induced E-fields in a FEM model of the human head.

We tested our pipeline on MRI-based models of the human cerebellum from five individuals [38] for two coil types, i.e., MagStim D70 and Deymed 120BFV [39], and we used lobule VIII as the primary ROI due to the potential application in movement disorders. Through computer simulations, we found that our algorithm predicts the cellular recruitment elicited by TMS pulses up to 600 A/µs with mean absolute percentual errors below 2%. We also found that, for TMS pulse intensities within the recommended ranges for human subjects [40], significantly more axonal fibers stemming from lobule VIII can be recruited by placing the coil at the optimal locations determined by our algorithm compared to position *3L1I*. Finally, we found that, by placing the coil at the proposed optimal positions, the minimum pulse intensity to recruit fibers on a 4 mm^2^-surface of the lobule VIII decreases by 21.57% (range: 4.68-55.05%) and 10.73% (range: 7.93-18.16%) (MagStim D70 and Deymed 120BFV, respectively) compared to *3L1I*, while the spillover to the adjacent lobule VII decreases by 70.61±16.27% and 71.74±20.83% (mean±S.D.), respectively.

Overall, our tool provides an effective solution to rapidly predict the acute response to TMS, inform parameter selection for cerebellar stimulation, and adjust the coil’s placement in patient-specific manner to maximize the cellular response to TMS.

## MATERIALS AND METHODS

Our multiscale computational pipeline was developed by integrating TMS-induced E-fields calculated in finite element models of the human cerebellum and multicompartmental Purkinje cell models endowed with realistic morphologies. The proposed algorithms were then developed and tested in this pipeline.

### MRI-based Atlases and Finite Element Model of the TMS-induced E-fields

High-resolution parcellations from 3T MRI scans of five healthy volunteers (both T1- and T2-weighted; resolution: 0.3 mm) were considered in this study. Parcellations were presented in [38] and include a manual parcellation of the cerebellar lobules and deep nuclei. Since the parcellation in [38] does not account for the vermis-hemispheric boundaries, the imaged cerebelli were further processed by using the ACAPULCO software [41] to retrieve the vermis parcels, ***Supplementary Information*, Fig. S1A-B**. Parceled cerebelli were imported in SimNIBS ver. 4.0 [42] for mesh generation and processed by using the CHARM pipeline to extract the head tissues [43]. Since the atlases were obtained from above-mouth scans, a neck section of 150 voxels along the z-axis was taken from [44] and automatically appended at the bottom of each image without altering the lobule segmentation prior to mesh generation. Nine tissue types were segmented, with each tissue type assigned a mean conductivity value as in [45, 46], ***Supplementary Information*, Table S1**.

For each head model, we considered several combinations of coil position and orientation (*see next sections*), and, for each combination, FEM calculations of the E-field were conducted in SimNIBS with models of the MagStim D70 (Magstim Inc., Whitland, UK) and Deymed 120BFV (Deymed, Hronov, Czech Republic) coils [39]. In all combinations, E-fields were modulated temporally with the biphasic TMS pulse waveform generated by the MagPro X100 stimulator (MagVenture A/S, Farum, Denmark; duration: 300 µs/phase; sampling rate: 5 MHz), and E-field distributions were calculated under quasi-static approximation [47] as proposed in [48], i.e., E-fields were first decomposed in a spatial component and a time component; then, the relative spatial distribution of the E-fields, which is time-invariant [49], was computed for a single coil current rate of change (1 A/µs; *normal* E-fields); finally, at each location, the E-field amplitude (in V/m) at any point in time was calculated as the product between the *normal* E-field and the coil current rate of change (in A/µs) at that point.

### Coupling E-fields to Neuron Models

The Purkinje layer was approximated by interpolating the surface mesh at the boundary between gray matter and white matter (2.81±0.82 million triangular elements, yielding a density of 35.19±0.60 elements per mm^2^, mean±S.D. across all five head models) and placing one Purkinje cell (PC) model per mesh point with the soma centered in the vertex and the somato-dendritic axis orthogonal to the outer surface. This was done because, even though the Purkinje layer lies in the gray matter and away from the boundary with the white matter [23], the median thickness of the granule layer underneath the Purkinje layer is less than the resolution of our MRI images [50]. Cells forming the granule layer (e.g., granule cells, Golgi cells, etc.) were omitted in this study because of their size, which is significantly smaller compared to Purkinje cells [51] and contributes to their low response to E-fields [52].

All PC models were implemented in NEURON ver. 8.0 [53] and modified from the multicompartmental cable model in [54] by adding a complete axon to the original morphology of soma, dendritic tree, and axon stem. The axon was attached after the second node of Ranvier (NoR) of the axon stem (i.e., 227 µm from the soma), which roughly corresponds to the region where sharp bends in the projecting axon are observed [29, 30]. Shape, orientation, and length of the axon varied with the mesh point where the PC model was placed and were determined by calculating the shortest path from the vertex to the volume representing the corpus medullare (CM). Briefly, for each PC model, we used the geodesic distance algorithm [55] to find the shortest path that the axon must transverse to connect the mesh vertex to CM, where the “shortest” path is the sequence of consecutive voxels that minimizes the quasi-Euclidean distance between the vertex and CM under the constraint of lying within the white matter and at least 16 voxels from any boundary with the gray matter. Then, we sampled and smoothed the shortest path via cubic spline interpolation, and we developed the axon along this path as an alternated sequence of myelinated compartments and NoR, **Fig. 1A**. Diameter, length, membrane properties, and ion channel composition of the myelinated compartments and the NoR were kept as in [54]. To account for the anisotropic arrangement of the axon terminations onto the deep cerebellar nuclei [29], a random rotation was applied to the distal portion of each axon, i.e., a point was randomly picked along each axon so that the distal segment includes the last 5-to-10 NoR, and this segment was rotated between 0° and 90° (uniform distribution) in any direction orthogonally to the plane of the proximal segment.

**Figure 1.**
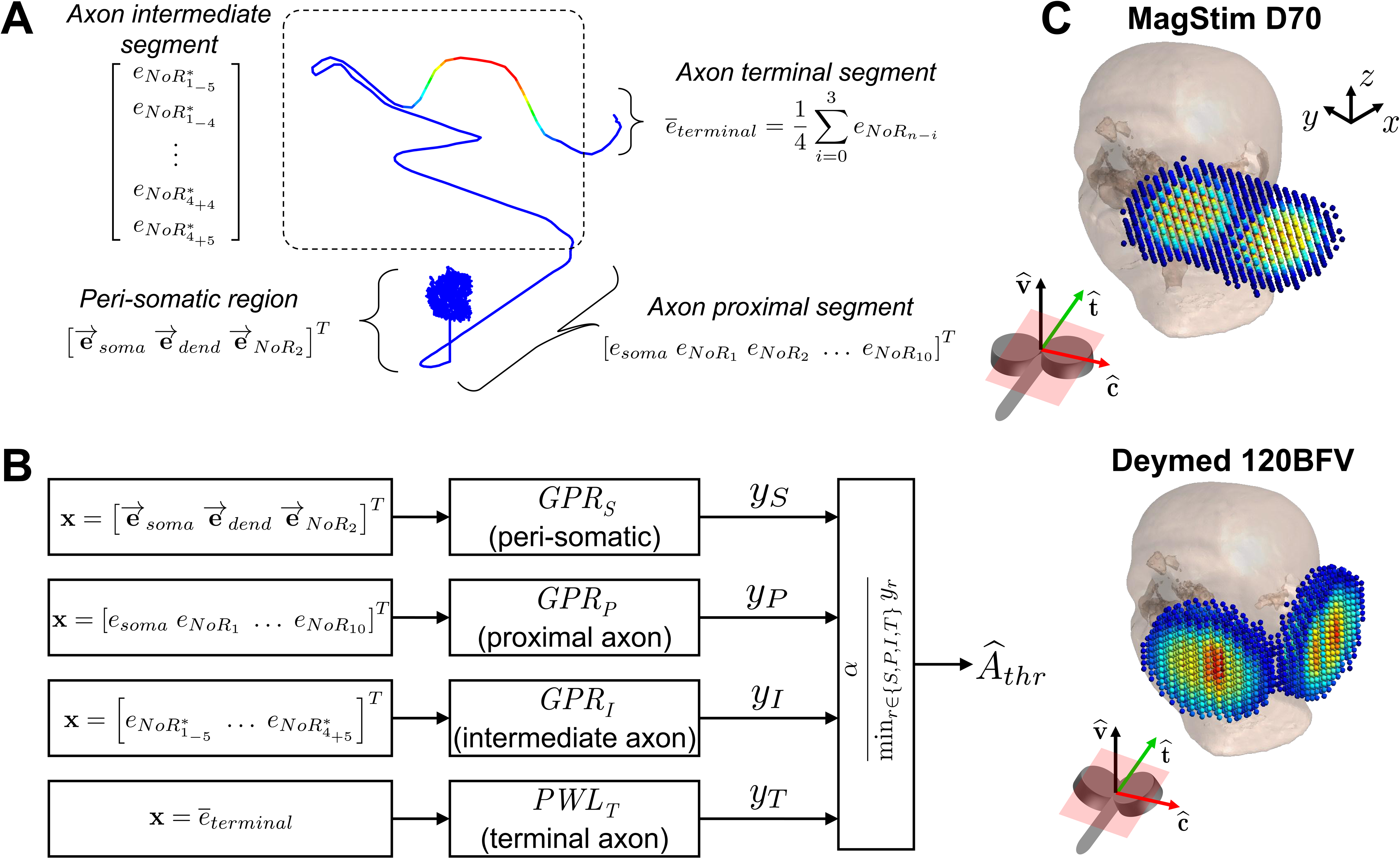
**A)** Segmentation of the PC model in partially overlapping regions and corresponding feature sets used to predict the region’s activation threshold. For the peri-somatic region, the local E-field 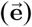 at the soma, distal dendrite (*dend*), and second node of Ranvier (*NOR*_2_) is calculated. For each compartment, 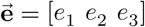 is a row vector including the E-field projections along the compartment’s longitudinal, transverse, and orthogonal axes, respectively. The feature set **x** for the peri-somatic region is a 9×1 vector. For the axon *proximal* segment, the feature set **x** is a 11×1 vector including the E-field projections at the soma and first 10 nodes of Ranvier (*NOR_i_*, *i* =1, 2, …, 10). For the axon *intermediate* segment, **x** is a 44×1 vector consisting of the E-field projections at the *critical* nodes of Ranvier 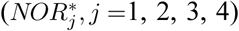 and, for each critical node, the 5 nodes of Ranvier preceding and the 5 nodes following the critical node (i.e., 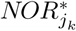, *k* = ±1, ±2, …, ±5). For the axon *terminal* segment, **x** is a scalar, i.e., the average E-field projection, *e̅_terminal_*, calculated across the last four nodes of Ranvier. **B)** Schematic of the algorithm for fast Purkinje cell activation threshold prediction. Feature sets calculated for every neuron region are fed to separate predictive models (GPR or PWL), and the minimum output value is used to provide the predicted activation threshold, *Â_thr_*. **C)** Magnetic vector potential as reconstructed in SimNIBS from the dipole expansion at the surface of the MagStim D70 coil (*top*) and Deymed 120BFV coil (*bottom*). For sake of visualization, coils are positioned on to the left cerebellar region in the MNI 152 head template. The MNI coordinate system (*x*, *y*, *z*) is also depicted. *Insets:* Unit vectors indicating the coil’s longitudinal axis (**ĉ**, *red*), transverse axis (**t̂**, *green*), and orthogonal axis (**v̂**, *black*). The plane defined by **ĉ** and **t̂** (*light red*) is also shown.

To couple the TMS-induced E-fields with the PC models, we used the generalized cable equation [56, 57]. First, we computed the spatial distribution of the *quasipotentials* associated with the E-field by numerically integrating the spatial component of the E-field along the compartmental sections of the PC models, with the soma being kept as reference (i.e., the *quasipotential* was 0 mV at the soma) [48, 57]. The E-field at each compartmental section was obtained via bilinear interpolation of the values at the 10 nearest mesh points. Then, the *quasipotentials* were applied to the center of the compartmental sections using the **`extracellular`** mechanism in NEURON [53]. Finally, the temporal component of the E-field was included by uniformly scaling the spatial distribution of the *quasipotentials* over time by the TMS pulse waveform. All neuron models under TMS-induced E-fields were discretized in compartmental sections no longer than 20 µm and simulated at a temperature of 37°C (CVODE solver; integration step: 0.005 ms) on the University of Connecticut High Performance Computing cluster.

### Algorithm for Fast Purkinje Cell Activation Threshold Prediction

In our coupled E-field/cerebellum models, Purkinje cells were “activated” if the membrane potential crossed −20 mV at the soma or 0 mV at any axonal compartment with a positive slope, as the presence of any of these two conditions was always followed by the initiation of an action potential. For any TMS-induced E-field, the “activation threshold” of a Purkinje cell was defined as the minimum coil current’s rate of change at the pulse onset, *A_thr_*, required to activate the PC model. Note that the activation threshold varies with (i) the relative spatial distribution of the E-field within the cerebellum, which depends on the specific brain atlas’ anatomy and the type, position, and orientation of the coil, and (ii) the location of the PC model within the cerebellar mesh, as this affects both the relative orientation of the PC model compared to the local E-field and the length and shape of the attached axon.

Because of the large number of combinations stemming from (i)-(ii), we developed a machine learning algorithm that predicts the activation threshold of the multicompartmental PC models without simulating these models. This algorithm combines Gaussian process regression (GPR) and piecewise linear (PWL) models and predicts the activation threshold of any PC by solely using the local E-field at the soma, distal dendrite, and NoR compartments, **Fig. 1B**.

First, the Purkinje cell is divided in four partially overlapping regions: (1) the peri-somatic region, which includes the soma, dendritic tree, and initial axon stem, i.e., up to the sharp bend in the axon; (2) the axon *proximal* segment, which spans the first 10 NoR from the soma and covers the axonal bend; (3) the axon *terminal* segment, which spans the last 4 NoR, i.e., the farthest nodes from the soma; and (4) the axon *intermediate* segment, i.e., the portion of axon between the proximal and terminal segments, **Fig. 1A**. Since the preferred site of action potential initiation in Purkinje cells is located around the first axonal branching point [32], these four neuron regions are defined to capture the different levels of sensitivity to the local E-field intensity and gradient that are expected along the membrane of Purkinje cells.

Secondly, for each neuron region, a set of features is defined to capture the sensitivity to the local E-field, **Fig. 1A**. It is hypothesized that the activation threshold for the *terminal* segment depends on the average value of the intensity (*e*) of the E-field projections along the nodes of Ranvier in the segment, i.e., the local E-field at each NoR is projected along the node’s longitudinal axis and averaged across all NoR in the segment. The local E-field at any compartment was the difference between *quasipotentials* at the compartment and its child compartment.

For the peri-somatic region, instead, the activation threshold is expected to depend on the entire E-field vector (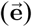) locally measured at the soma compartment, the axonal bending site (i.e., the second NoR), and the most distal dendritic compartment (i.e., 305.5 μm from the soma). For the *proximal* segment, the activation threshold is expected to depend on the intensity of the E-field projections around the bending site, i.e., from the soma compartment to the 10^th^ node of Ranvier. Finally, for the axon *intermediate* segment, we accounted for the fact that the number of NoR vary with the specific PC model, and therefore we considered only four *critical* nodes (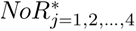*NoR*^∗^, **Fig. 1A**). Briefly, we calculated the E-field projection and gradient along the longitudinal axes of the nodes of Ranvier and defined the *critical* nodes as the four

NoR at which either the E-field projection intensity or the E-field gradient reach the most positive value or the most negative value (i.e., minimum). For every *critical* NoR, we then consider the intensity of the E-field projections at the closest five NoR preceding the node and the five NoR following the node, thus gathering a feature vector that includes 11 intensity values of E-field projections per *critical* NoR.

Finally, the E-field values considered for each neuron region are fed to separate predictive models, one per region, **Fig. 1B**. Denoted with **x** the set of E-field values (*feature set*) for any of the four neuron regions, the model’s output is defined as follows:

- *y_r_* = [1 **x***^T^*]**b***_r_* + *f_r_*(**x**), in case of peri-somatic region (*r* = *S*), axon *proximal* segment (*r* = *P*), or axon *intermediate* segment (*r* = *I*), where **x** is a column vector.
- *y_r_* = *k_r_*(**x**)**x**, in case of the axon *terminal* segment (*r* = *T*), where **x** is a scalar.

The neuron’s predicted activation threshold is provided as *Â_thr_* = *⍺*⁄*y_r_*, where *⍺* = 10^3^ is a scaling factor. For *r* = *T*, a PWL predictive model is used, i.e., *k_r_*(*x*) = *k_l_* if *x* ≥ 0 and *k_r_*(*x*) = *k_2_* if *x* < 0. For *r* = *S*, *P*, *I*, a GPR predictive model is used, where **b***_r_* is a parameter vector to be estimated, and 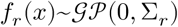 is a Gaussian process with zero-mean and covariance function, Σ*_r_*, given by exponential kernels, i.e., 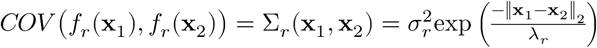 for any vectors **x**_1_ and **x**_2_.

We estimated the parameters 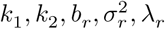 for *r* = *S, P, I*, and tested the prediction algorithm on the activation thresholds of a subset of PC models uniformly distributed across the left cerebellar hemisphere. Namely, for each head model, we placed the Deymed 120BFV coil as calculated by our optimization routine (i.e., in the positions “*L8p*” defined in the next section; see coordinates in ***Supplementary Information*, Table S2**) and calculated the *normal* E-fields. Then, we paired the *normal* E-fields with 40,000 PC models (200,000 across all head models) randomly distributed across the left lobules V, VI, VII, and VIII. Finally, for each PC model, the coil current rate of change was scaled according to a binary search algorithm until the minimum intensity, *A_thr_*, necessary to elicit an action potential was achieved with 0.5 A/µs accuracy. See text in ***Supplementary Information*** for details on the training and testing of the prediction algorithm.

All computations were conducted in MATLAB, rel. R2021b, The MathWorks, Inc., Natick, MA, on a Fedora Linux 35 twelve-core Intel Xeon workstation (3.40 GHz/core, 50 GB RAM).

### Algorithm for Fast Optimization of Coil Placement

We developed a multistep algorithm for fast calculation of the coil’s position and orientation maximizing the TMS-induced E-field in a desired cerebellar ROI. Albeit applicable to any ROI, we targeted the left lobule VIII because of its involvement in motor functions [58, 59]. The centroid of the mesh points spanning the left lobule VIII parcel, including the subregions VIII-A and VIII-B, was used as the fiducial reference point, *c_ref_*, and a minimum distance of 2 mm between any point of the coil and the mesh grid of the outer surface of the head model (i.e., scalp) was always enforced. The following steps were repeated for every combination of coil type and head model under the assumption that coils have axial symmetry (i.e., we do not control the coil current direction), and the E-fields through the brain have equal magnitudes and reverse directions when the direction of the coil’s current is flipped [48].

- ***Step 1* (Initialization):** A sphere centered in the fiducial reference, *c_ref_*, is drawn with radius gradually increased until the intersection with the scalp mesh grid includes at least 30 mesh points (*candidate* points). The minimum number of *candidate* points was set to provide a dense grid of positions at the scalp around the fiducial reference point. For each *candidate* point, the coil is placed with the face toward the scalp, and the coil center is moved at least 30 mm (MagStim D70: 30 mm; Deymed 120BFV: half the coil’s depth) from the *candidate* point along the direction of the normal vector, **n̂**, to the scalp’s outer surface, while the coil’s orthogonal axis (unit vector **v̂** in **Fig. 1C**, *Insets*) is aligned with **n̂**. A total of 36 initial positions per *candidate* point are generated by rotating the coil around the direction of **n̂**, i.e., the orientation of the coil is varied from −90° to +80° (increments of 10°) along the transversal plane (*light red* in **Fig. 1C**, *Insets*).
- ***Step 2* (Coil’s Surface-to-Scalp Projection Maximization):** The coil is placed in every initial position indicated at *Step 1* and rotated around the unit vectors **t̂** (i.e., along the plane defined by **v̂** and **ĉ**) and **ĉ** (i.e., along the plane defined by **v̂** and **t̂**), **Fig. 1C**, to enhance the projected coil surface onto the scalp. For every initial position, *w*, the best rotation angles 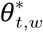 and 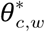 around **t̂** and **ĉ**, respectively, are determined as the solution to the constrained minimization problem:

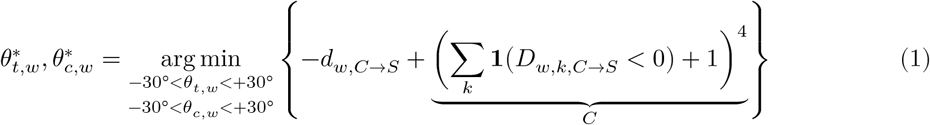 where *θ_t,w_* and *θ_c,w_* indicate the rotation angles around **t̂** and **ĉ**, respectively, when the coil is in the initial position *w*, *D_w,k,C→S_* is the Euclidean distance between any mesh point *k* of the coil model in position *w* and the closest mesh point (i.e., node) on the scalp grid, and *d_w,C→S_* is the minimum distance between coil and scalp, i.e., 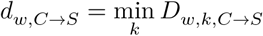. Note that the distances *D_w,k,C→S_* depend on *θ_t,w_* and *θ_c,w_*, and the step function **1**(*D_w,k,C→S_* < 0) is positive if the coil’s mesh point *k* moves underneath the scalp surface because of the rotations and is zero otherwise. Hence, the penalty term *C* in (1) aims to avoid rotations that would move the coil underneath the scalp. Also, −*d_w,C→S_* in (1) decreases as *d_w,C→S_* increases, which means that the solution to (1) maximizes the minimum distance between coil and scalp. Angles 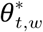 and 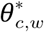 are calculated by using the **`fmincon`** method in MATLAB and result in maximizing the projected coil surface onto the scalp while the coil is in position *w*.
- ***Step 3* (Coil’s Center-to-Scalp Distance Minimization):** Starting from the site defined by the initial position *w* and the rotation 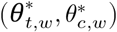 at the end of *Step 2*, the coil is translated along the direction of **n̂** until a new location *w_l_*, with 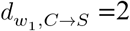 mm, is reached. Then, position and orientation of the coil are adjusted to further enhance the projected coil surface on the scalp while keeping the coil-to-scalp distance as close as possible to 2 mm. This is obtained by minimizing the distance between the coil’s center and the scalp while preventing that any coil’s mesh point moves closer than 2 mm from the scalp, i.e., the best rotation angles 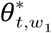 and 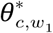 around **t̂** and **ĉ**, respectively, and the best advancement, 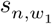, (in mm) of the coil’s center away from the location *w*_l_ and along the direction of **n̂** are determined by solving the constrained minimization problem:

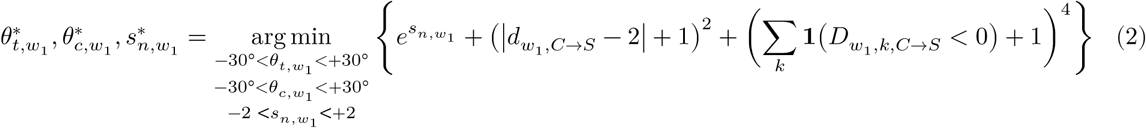 The solution to (2) is calculated by using the **fmincon** method in MATLAB to maximize the projected coil surface on the scalp while the coil is around position *w*_l_ and the minimum coil-to-scalp distance is 2 mm.

For any initial position *w*, *Step 1-3* provide a final location (position and orientation) of the coil. The *normal* E-field is calculated for every final location, and the optimal placement, which is labeled “*L8p*”, is given by the location with the highest average *normal* E-field intensity in the ROI.

## RESULTS

We tested our algorithms on five head models with cerebellar lobule parcellation from [38]. In each model, Purkinje cells were placed in the nodes of the mesh grid spanning the Purkinje layer and initialized at −60 mV. To focus on the effects of E-fields on the Purkinje cell’s firing, PC models were maintained silent at rest. Also, PC models were augmented by adding axons with 3D geometry that extend through the white matter from the Purkinje layer to the CM. The arrangement of the axonal fibers (length: 54.37± 22.07 mm, mean±S.D.) was characterized by long, nonplanar pathways running in the posteroanterior direction. These pathways started with initial segregation in the cerebellar lobules and rapidly formed large and densely organized bundles towards the CM, **Fig. 2A**. Moreover, although the distribution of axonal pathways varied with the size, shape, and parcellation of the cerebellar volume, ***Supplementary Information*, Fig. S2A-D**, all fiber bundles presented limited symmetry across hemispheres, which enabled a realistic investigation of the spillover effects of TMS on the right hemisphere when the coil is placed on the left side.

**Figure 2.**
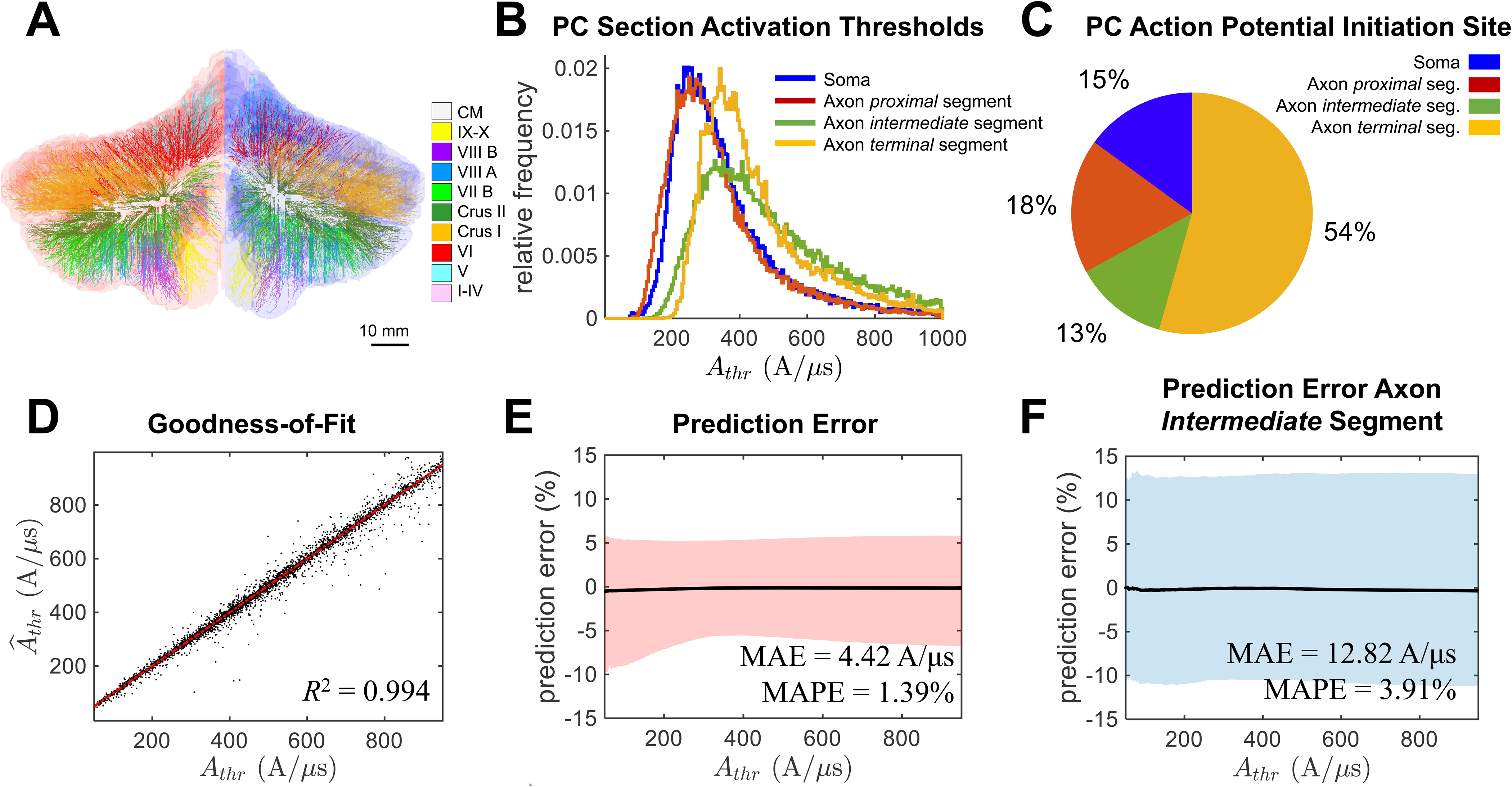
**A)** 3D arrangement of the multicompartmental axons attached to the Purkinje cell models in one head model (Atlas 2). Note that only 1% of the estimated axons are depicted. Axons are colored according to the legend on the right to indicate the lobules from which they depart (CM: corpus medullare; I-X: lobules I-X, including the sub-lobule regions Crus I, Crus II, VII-B, VIII-A, and VIII-B). **B)** Probability distribution function across five head models of the minimum TMS pulse intensities (i.e., activation thresholds) at which an action potential is initiated at the soma, axon *proximal* segment, axon *intermediate* segment, or axon *terminal* segment of the simulated Purkinje cell models. **C)** Percentage of simulated Purkinje cell models across five head models for which an action potential initiates at the soma, *proximal* segment, axon *intermediate* segment, or axon *terminal* segment when the activation threshold is reached. **D)** Activation threshold, *Â_thr_*, predicted by the algorithm in Fig. 1B versus the actual activation threshold, *A_thr_*. Black dots denote Purkinje cell models simulated in Atlas 1, 3, 4, and 5 (*test data*) while the algorithm was trained on data from Atlas 2. A linear regressor (*red* line) is fit on the black dots, and the goodness-of-fit is provided by the coefficient of determination, *R*^2^. **E)** Prediction error calculated for the Purkinje cell models in **D)** and sorted according to increasing values of the actual threshold, *A_thr_*. **F)** Prediction error calculated for the Purkinje cell models in **D)** whose activation occurs at the axon *intermediate* segment. In **E-F)**, for each value of *A_thr_*, the prediction error is expressed as percentage, i.e., 100 × (*Â_thr_* − *A_thr_*)⁄*A_thr_*. For each value of *A_thr_*, the prediction error is reported as mean (*black* line) and 95^th^ percentile region (colored region). The mean absolute error (MAE) and mean absolute percentage error (MAPE) across all values *A_thr_* are reported.

### GPR Models Predict Purkinje Cell Activation Thresholds with High Accuracy

A total of 200,000 PC models (40,000 per head model) randomly distributed across the left lobule V, VI, VII, and VIII were simulated, and the corresponding activation thresholds were used to test the algorithm for fast PC activation threshold prediction.

Across all computer simulations, we found that the activation thresholds were generally higher than the pulse intensities typically used in TMS studies and concentrated between 150 A/μs and 600 A/μs, **Fig. 2B**, which suggests that direct activation of the Purkinje layer may be limited. Furthermore, even though the average intensity required to initiate an action potential at the PC soma or axon *proximal* segment was lower compared to the distal axon segments (370.1 A/μs, 353.9 A/μs, 533.8 A/μs, and 473.6 A/μs for the PC soma and axon *proximal*, *intermediate*, and *terminal* segment, respectively; one-way ANOVA test, *P*-value *P*<0.05), less than 30% of the Purkinje cells across lobule V-VIII would initiate an action potential at the soma or the axon *proximal* segment when the coil is in *L8p*, **Fig. 2C**. Also, the Purkinje cells with low somatic activation thresholds were mostly located on mesh points on gyral crowns directly in front of the coil, which suggests that the somato-dendritic axes of these neuron models were closely aligned with the normal component of the E-fields. Finally, a preference for activating PC axon terminals was found, **Fig. 2C**, which is consistent with the effects of cerebral TMS on pyramidal neurons [48] and is further supported by experimental data on corticospinal activation [60, 61].

A five-fold cross-validation scheme was used to assess our prediction algorithm, i.e., at each fold, the predictive models in **Fig. 1B** were fitted on data from one head model, and the quality of the prediction was measured on data from the remaining four head models (*Test Set*) in terms of mean absolute error (MAE) and mean absolute percentage error (MAPE). **Fig. 2D-F** show the performance for one *Test Set* (i.e., models were trained on data from Atlas 2).

First, we noticed that the training was fast (i.e., less than 5 minutes per fold), mainly because of the small number of model parameters to estimate. Also, when applied to test data, the algorithm resulted in a strong correlation between predictions, *Â_thr_*, and corresponding true values, *A_thr_*, **Fig. 2D** (coefficient of determination, *R*^2^>0.99), which remained consistent across *Test Sets*, ***Supplementary Information*, Table S3** (last row). Also, both GPR models and PWL models provided predictions that are linearly related to the simulated thresholds, see ***Supplementary Information*, Fig. S3A-D** and **Table S3**, thus indicating that the proposed E-field-based features are highly correlated with changes to transmembrane voltages along the Purkinje cells.

Secondly, we considered the prediction error versus simulated thresholds, **Fig. 2E**, and found that the average MAE was low, i.e., 4.91±0.49 A/µs (mean±S.D. across five *Test Sets*), remained consistent across *Test Sets* (see ***Supplementary Information*, Table S4**), and resulted in relative errors below 2% (MAPE: 1.61±0.20%, mean±S.D. across five *Test Sets*). Finally, even though GPR and PWL models had similar predictive power, the prediction error was generally higher for the axon *intermediate* segment compared to the other segments, **Fig. 2F** and ***Supplementary Information*, Table S4**, which is likely due to the variety of shapes and 3D pathways encompassed by the axon *intermediate* segments compared to the *proximal* and *terminal* segments. Nonetheless, the MAPE always remained below 5%.

Given the postero-anterior arrangement of the Purkinje cells along the cerebellar gyri, we tested whether the E-field at the soma could be a simpler, yet effective proxy to predict the activation of Purkinje cells. We considered the E-field projection (*e_soma_*) along the somato-axonal axis of the simulated Purkinje cells and calculated the linear fit between *e_soma_* and the outcome, *y_S_*, of the somatic GPR model and between *e_soma_* and the reciprocal of the prediction *Â_thr_*, ***Supplementary Information*, Fig. S4A-B**. We found that, albeit correlated with *y_S_*, *e_soma_* is a poor predictor of the cell’s overall activation threshold. Similarly, the E-field magnitude at the soma, 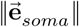, was a poor predictor of both *y_S_* and the reciprocal of *Â_thr_*, even though it improved compared to the projection *e_soma_*, ***Supplementary Information*, Fig. S4C-D**. Moreover, 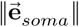 was a weak predictor of the activation threshold of each axonal segment considered in our study, i.e., *y_P_*, *y_I_*, and *y_T_*, ***Supplementary Information*, Fig. S5A-C**.

Altogether, our results indicate that, by exploiting features that capture the spatial distribution of the E-field along the axonal fibers, our algorithm can predict the activation threshold of Purkinje cells with low absolute error. The algorithm is robust against wide fluctuations in the shape, length, and orientation of the PC axon, can be trained quickly, and significantly improves over the predictions based on the sole magnitude of the E-fields along the cerebellar gyri, which is consistent with experimental data in [62].

### Optimized Coil Placement Increases the E-field Intensity in the ROI

*Step 1-3* of the coil placement optimization algorithm were repeated for 1,070±418 (MagStim D70) and 533±85 (Deymed 120BFV) *candidate* combinations of coil positions and orientations per head model (mean±S.D. across five head models). The optimal placement (*L8p*) was selected as the combination that maximizes the average *normal* E-field intensity in the left lobule VIII (ROI). All results are reported as mean±S.D. across five head models.

We found that the location *L8p* varied across head models and coil types. However, upon projecting the *L8p* positions calculated for every head model on the MNI 152 template [63], we found that the Euclidean distances between these locations were small for both coil types, see **Fig. 3A** (***i***) and **Fig. 3B** (***i***) for MagStim D70 and Deymed 120BFV, respectively. By comparing the *L8p* locations across coil types (***Supplementary Information*, Table S2**), we found that the Deymed 120BFV coil had locations more lateral and elevated away from the scalp compared to the MagStim D70 coil (71.32±6.81 mm vs. 63.70±5.05 mm lateral and 26.70±8.94 mm vs. 23.65±2.87 mm inferior to the inion; average distance of the coil center from the scalp: 14.89±2.92 mm vs. 3.96±1.04 mm; minimum distance of the coil center from the cerebellar grey matter surface: 29.65±3.67 mm vs. 16.68±1.70 mm; Deymed 120BFV vs. MagStim D70), **Fig. 3A-B** (***ii***). Perhaps more importantly, the Deymed 120BFV coil surface was more directly facing the scalp compared to the MagStim D70 coil, i.e., the angle, *φ*, between the *y*-axis in **Fig. 3A-B** (***i***) and the coil’s unit vector **v̂** was 11.03°±3.96° vs. 31.90°±12.56° (Deymed 120BFV vs. MagStim D70).

**Figure 3.**
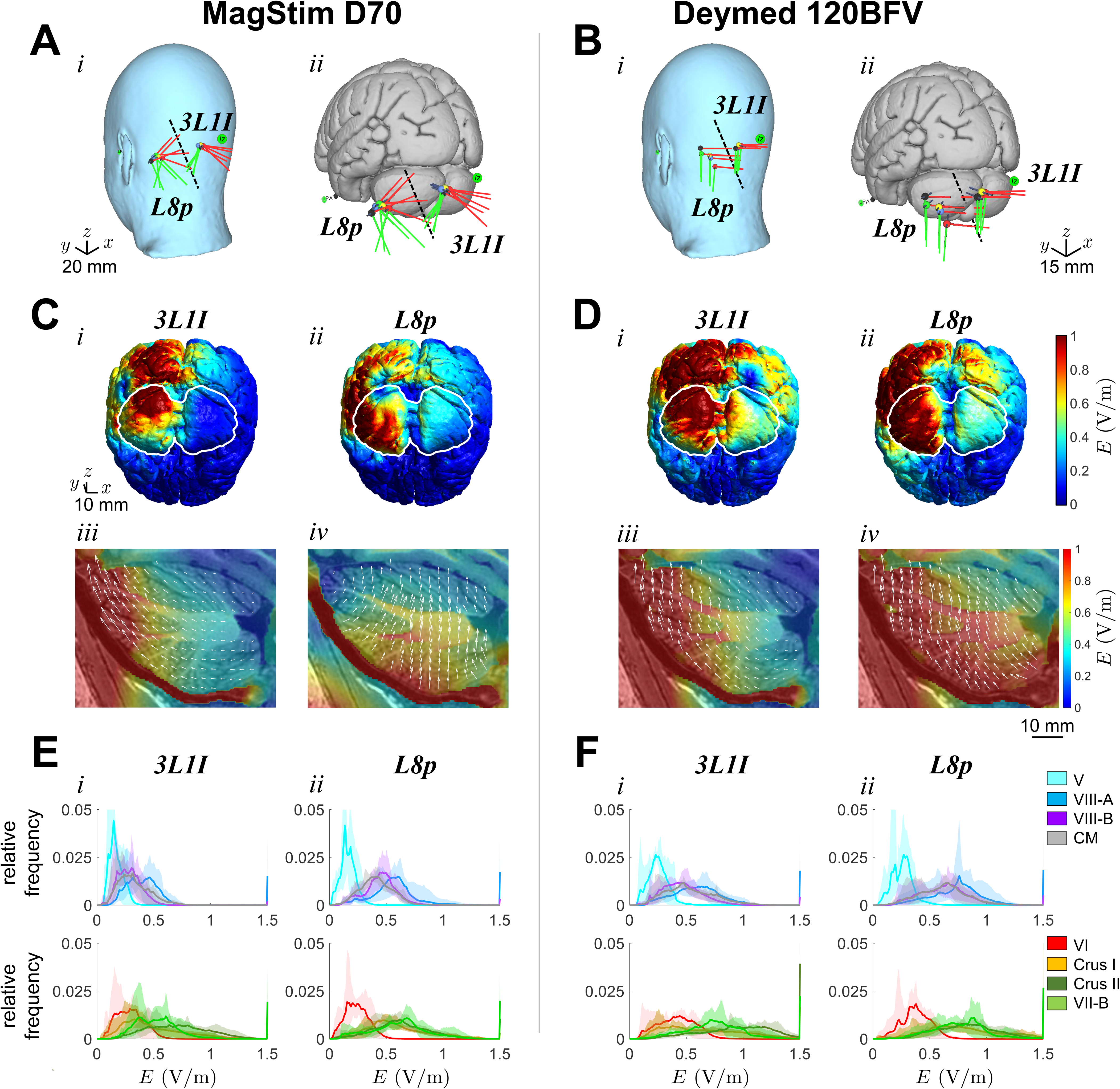
Optimal coil placement (*L8p*) and position *3L1I* in all head models and corresponding *normal* E-field distribution calculated for the MagStim D70 coil (**A**, **C**, **E**) and Deymed 120BFV coil (**B**, **D**, **F**). **A-B)** Projection of the calculated positions *L8p* and *3L1I* on the MNI152 atlas when scalp layers are added (***i***) and the gray matter is exposed (***ii***), respectively. In each panel, the colored balls indicate the coil’s center point for five head models (*yellow*: Atlas 1; *blue*: Atlas 2; *red*: Atlas 3; *green*: Atlas 4; *black*: Atlas 5), and the lines indicate the orientation of the coil’s longitudinal and transverse axes, i.e., **ĉ** (*red lines*) and **t̂** (*green lines*), respectively. The scales in **A)** (bottom left) and **B)** (bottom right) apply to panels (***i***) and panels (***ii***), respectively. **C-D)** *Normal* E-field across the cerebellar gray matter in one head model (Atlas 5) for the TMS coil applied in position *3L1I* (***i***, ***iii***) or *L8p* (***ii***, ***iv***) with coil current flowing downwards. For each coil position, a posterior-inferior view of the E-field on the gray matter surface (***i***-***ii***) and a sagittal slice of the left cerebellar hemisphere (***iii***-***iv***) are shown. The boundary of the cerebellar region in (***i***-***ii***) is marked in white. The scale in **C-*i*)** applies to **C-D)**, panels (***i***-***ii***). The scale in **D-*iv*)** applies to **C-D)**, panels (***iii***-***iv***). The color maps in **D**-***ii*)** and **D**-***iv*)** apply to panels (***i*-*ii***) and panels (***iii*-*iv***), respectively, in **C-D)**. **E)** Sample distribution of the *normal* E-field intensities within each cerebellar lobule when the coil is in position *3L1I* (***i***) or *L8p* (***ii***). Distributions are shown separately for lobules that are primarily invested in motor functions (*top row*) and non-motor functions (*bottom row*). Distributions are computed separately for each head model and reported as mean (*solid line*) and min-max range across the head models (*shaded area*).

To mimic the procedures followed in experimental settings to place the coil 3 cm lateral and 1 cm inferior to the inion, we consider a fiducial marker 3 cm lateral and 1 cm inferior to the inion in every head model, and we applied *Step 1-3* to this marker, i.e., the marker was treated as a *candidate* point. The resultant adjusted positions are labeled “*3L1I*”, and the corresponding coordinates are reported in ***Supplementary Information*, Table S2** for all head models. Positions *3L1I* aimed to tune the coil orientation and distance from the scalp around the fiducial marker and strengthen the cerebellar response to E-fields.

Positions *3L1I* resulted in a more central and elevated placement of the coil compared to *L8p* (*3L1I*: 27.93±2.03 mm lateral and 9.89±1.71 mm inferior to the inion, *φ* =25.77°±12.49° for MagStim D70, and 35.20±3.65 mm lateral and 13.42±2.16 mm inferior to the inion, *φ* =10.99°±3.93° for Deymed 120BFV; *L8p*: 63.69±5.05 mm lateral and 23.19± 2.87 mm inferior to the inion, *φ* =32.05°±12.42° for MagStim D70, and 71.32±6.81 mm lateral and 26.24±8.94 mm inferior to the inion, *φ* =11.15°±5.41° for the Deymed 120BFV coil). Also, when comparing *L8p* vs. *3L1I*, we found that, for both coil types, the average distance of the coil center from the ROI’s centroid and outer surface was significantly lower for coils in position *L8p* (Wilcoxon rank sum tests *L8p* vs. *3L1I*; *P*-value *P*<0.01) while the average distance from the surrounding lobules (i.e., left lobule V-VII) was similar for both positions, see ***Supplementary Information*, Fig. S6**.

The displacement between positions *L8p* and *3L1I* had remarkable effects on the spatial distribution of the E-fields across the cerebellar lobules for both coil types. **Fig. 3C-D** (***i***) and **Fig. 3C-D** (***ii***) show the calculated *normal* E-field in one head model (Atlas 5) when the coil is placed in positions *L8p* and *3L1I*, respectively. With both coils, positions *3L1I* led to high E-field intensities outside the cerebellum (white lines) and mainly over the primary visual cortex. Moreover, Deymed 120BFV coils caused a significant spillover to the right hemisphere, both in the cerebellum and occipital lobe, **Fig. 3D** (***i***). Vice versa, high E-field intensities were centered in the target cerebellar hemisphere for coils in *L8p*, and the spillover outside the cerebellum was mostly lateralized towards the mastoidal area, **Fig. 3C-D** (***ii***). Finally, positions *3L1I* generally resulted in lower E-field intensities in the target ROI compared to the surrounding lobules (i.e., lobule VII-B, Crus I, and Crus II), **Fig. 3C-D** (***iii***), mainly because of the inward orientation of lobule VIII. Positions *L8p*, instead, guaranteed higher E-field intensity in lobule VIII compared to *3L1I* and resulted in a general shift of the E-field distribution from the Crus area in lobule VII to lobule VIII, with the shift being consistent across head models (see **Table 1**) and more clearly observable for MagStim D70 coils compared to Deymed 120BFV coils, see **Fig. 3C** (***iv***) vs. **Fig. 3D** (***iv***).

**Table 1.** Average intensity of the *normal* E-field (in V/m) across five head models in the left cerebellar lobules V, VI, VII (including Crus I, Crus II, and VII-B), and VIII (including VIII-A and VIII-B), and corpus medullare (CM) for different coil types and locations. Average intensity is reported as mean (S.D.).

| Coil Type | Location | *Normal* E-field Intensity (V/m) in Left Cerebellum |  |  |  |  |  |  |  |
| --- | --- | --- | --- | --- | --- | --- | --- | --- | --- |
|  |  | V | VI | Crus I | Crus II | VII-B | VIII-A | VIII-B | CM |
| MagStim D70 | *3LII* | 0.169<br>(0.048) | 0.295<br>(0.099) | 0.498<br>(0.249) | 0.745<br>(0.432) | 0.597<br>(0.607) | 0.439<br>(0.350) | 0.292<br>(0.123) | 0.327<br>(0.121) |
|  | *L8p* | 0.162<br>(0.056) | 0.240<br>(0.085) | 0.591<br>(0.268) | 0.651<br>(0.407) | 0.702<br>(0.745) | 0.615<br>(0.507) | 0.454<br>(0.166) | 0.413<br>(0.138) |
| Deymed 120BFV | *3LII* | 0.250<br>(0.072) | 0.419<br>(0.141) | 0.665<br>(0.339) | 1.012<br>(0.705) | 0.873<br>(1.083) | 0.663<br>(0.633) | 0.476<br>(0.222) | 0.533<br>(0.195) |
|  | *L8p* | 0.258<br>(0.070) | 0.382<br>(0.103) | 0.808<br>(0.315) | 0.892<br>(0.707) | 0.938<br>(1.282) | 0.830<br>(0.818) | 0.636<br>(0.270) | 0.619<br>(0.177) |

Finally, we calculated the distribution of the *normal* E-field intensity separately for lobules V-VIII across all head models, both for coils in positions *3L1I* and *L8p*, **Fig. 3E-F** (***i***) and **Fig. 3E-F** (***ii***), respectively. We found that the transition from *3L1I* to *L8p* led to a significant increase in E-field intensity in motor-related lobules, i.e., lobules V, VIII-A, and VIII-B (top rows in **Fig. 3E-F**) for both coil types while the change remained modest for lobules involved in cognitive functions, especially lobule VI and Crus II (bottom rows in **Fig. 3E-F**). The cerebellar nuclei (i.e., CM in **Table 1** and **Fig. 3E-F**), instead, received E-field intensities comparable to the inner part of lobule VIII (i.e., VIII-B) and significantly lower compared to lobule VIII-A (paired *t*-test, *P*-value *P*<0.001).

Altogether, our optimization procedure led to new coil positions (i.e., *L8p*) that significantly increase E-field intensity in the cerebellar motor area, especially lobule VIII-A, compared to standard positions *L31I* while shifting the focus of the E-field distribution from the outer side of lobule VII (i.e., Crus I-II) to deeper cerebellar regions, i.e., lobule VII-B and VIII-A. This trend was more clearly observed for MagStim D70 coils because of the weaker, less focused E-fields, while Deymed 120BFV coils resulted in higher E-field intensities across most of the left cerebellar hemisphere.

### Predicted Purkinje Cell Activation for Optimal Coil Placement

We used our activation threshold prediction algorithm to calculate the intensity that TMS pulses should reach to activate Purkinje cells across the various cerebellar lobules for both coil types. This aimed to assess whether the optimized coil placement *L8p* can result in stronger neuronal response in the motor cerebellum compared to positions *3L1I*. For each head model, we used the trained prediction algorithm (i.e., the training was conducted on the other head models) to calculate the activation threshold at the nodes of the Purkinje layer mesh grid. Then, for each mesh triangle, the average predicted activation threshold for the Purkinje cells spanning the triangle’s surface was calculated as linear interpolation of the predictions at the vertices. **Fig. 4A-B** (***i***) and **Fig. 4A-B** (***ii***) show the progressive activation of the cerebellar lobules as the TMS pulse intensity rises (i.e., *activation map*) when the coil is in *3L1I* and *L8p*, respectively, for one head model.

**Figure 4.**
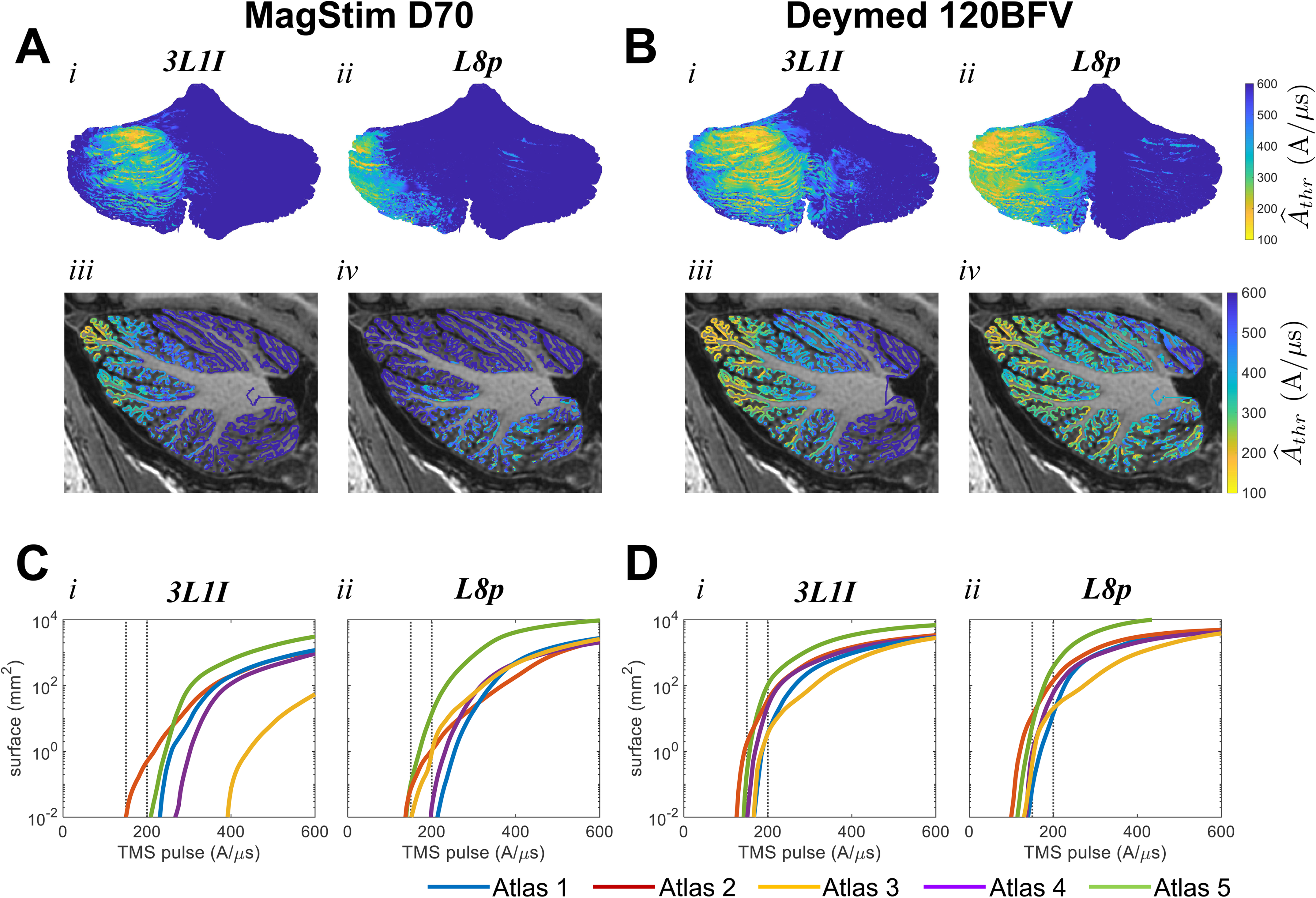
Predicted response of the Purkinje cells to a single TMS pulse when the MagStim D70 coil (**A**, **C**) or Deymed 120BFV coil (**B**, **D**) is used with coil current flowing downwards. **A-B)** Predicted activation thresholds, *Â_thr_*, calculated for the Purkinje cells in one head model (Atlas 5) for coil applied in position *3L1I* (***i***, ***iii***) or *L8p* (***ii***, ***iv***). For each position, a posterior view of the activation thresholds on the gray matter surface (***i***-***ii***) and a sagittal slice of the left cerebellar hemisphere with the activation thresholds (***iii***-***iv***) are shown. Color maps in **B-*ii*)** and **B-*iv*)** also apply to panels (***i***) and panels (***iii***), respectively, in **A-B)**. **C-D)** Total surface area (in mm^2^) activated by a TMS pulse in the left lobule VIII (i.e., VIII-A and VIII-B are combined) versus the pulse intensity (in A/μs) in all head models (Atlas 1-5). Surface scales in **C-D)**, panel (***i***) also apply to **C-D)**, panel (***ii***), respectively. In all panels, vertical *dotted* lines denote the total surface area activated by TMS pulses at 150 A/μs and 200 A/μs.

The activation maps for the MagStim D70 (**Fig. 4A**) and Deymed 120BFV (**Fig. 4B**) coils showed good alignment with the *normal* E-field distributions in **Fig. 3C** and **Fig. 3D**, respectively, i.e., the Purkinje cells with the lowest activation thresholds were located directly under the coil center, which also represented the region with the highest *normal* E-field intensities. This was particularly relevant for the activation of deeper regions, e.g., lobule VII-B, VIII-A, and VIII-B, which resulted in lower activation thresholds as the coil moved caudally from *3L1I* to *L8p*, see sagittal view in **Fig. 4A** (***iii***-***iv***) and **Fig. 4B** (***iii***-***iv***), respectively. Also, coils in position *3L1I* resulted in low-threshold surfaces mainly concentrated in the medial zone, i.e., the spinocerebellar area, while TMS pulses delivered at positions *L8p* primarily recruited neurons in the cerebrocerebellar area, **Fig. 4A-B** (***i***, ***iii***) vs. **Fig. 4A-B** (***ii***, ***iv***). Finally, the average activation threshold was lower for Deymed 120BFV coils compared to MagStim D70 coils and resulted in a larger area of activation for any given pulse intensity, **Fig. 4A** (***ii***) vs. **Fig. 4B** (***ii***).

To further quantify the neuronal response to cerebellar TMS, we varied the intensity of the TMS pulses, and, for each intensity, we calculated the portion of the mesh grid (i.e., surface area) that is activated by the pulses. We found that a 200-A/μs pulse delivered via MagStim D70 coil in position *3L1I* has limited impact on lobules VIII-A and VIII-B (a combined activated area was reported in just 2 out of 5 head models). The same pulse with the coil in position *L8p*, instead, resulted in a combined activated area in all head models (median: 0.90 mm^2^; range: [0.0004, 14.02] mm^2^). Similarly, the Deymed 120BFV coils resulted in larger activated area in lobules VIII-A and VIII-B for *L8p* compared to *3L1I* (median: 60.44 vs. 24.06 mm^2^; range: [14.00, 371.41] mm^2^ vs. [3.52, 106.00] mm^2^; *L8p* vs. *3L1I*; *P*-value *P*<0.001) and guaranteed better overall penetration compared to the MagStim D70 coils. Although these values are smaller compared to the values predicted for pyramidal neurons in cerebral TMS [48, 64], a monotonic trend was observed for all head models as the pulse intensity increases, with a consistent preference for positions *L8p* compared to *3L1I* for both coil types, **Fig. 4C-D** (***i***) vs. **Fig. 4C-D** (***ii***).

Finally, as for TMS in nonhuman primates [62], we calculated the minimum pulse intensity to activate an equivalent surface area of 1 mm^2^ (**Table 2**) or 4 mm^2^ (**Table 3**) of the left lobule VIII for every coil type and position. On average, the pulse intensity required to activate 4 mm^2^ was 8.85%±3.11% higher compared to the intensity that activates 1 mm^2^ (mean±S.D. across all head models, coil types, and positions). For MagStim D70 coils, positions *L8p* reduced the average intensity to activate 1 mm^2^ and 4 mm^2^ of lobule VIII by 23.20% (range: 1.34-54.43%) and 21.57% (range: 4.68-55.05%) (median across all head models), respectively, compared to *3L1I*, while the reduction was smaller and more uniform across head models for Deymed 120BFV coils (i.e., 13.11±3.29% and 12.76±4.87%, respectively).

**Table 2.** TMS pulse intensity (in A/μs) to activate a minimum surface of 1 mm^2^ of left lobule VIII (i.e., VIII-A and VIII-B combined) in each head model (i.e., Atlas 1-5) for different coil types and locations.

| Coil Type | Location | TMS pulse intensity (*A/μs*) to activate 1 mm<sup>2</sup> surface in left lobule VIII |  |  |  |  |
| --- | --- | --- | --- | --- | --- | --- |
|  |  | Atlas 1 | Atlas 2 | Atlas 3 | Atlas 4 | Atlas 5 |
| MagStim D70 | *3LII* | 257.85 | 217.09 | 440.45 | 302.17 | 243.92 |
|  | *L8p* | 254.40 | 197.92 | 200.71 | 232.08 | 170.68 |
| Deymed 120BFV | *3LII* | 186.37 | 144.07 | 185.11 | 169.71 | 156.46 |
|  | *L8p* | 169.43 | 120.48 | 154.32 | 150.36 | 137.58 |

**Table 3.** TMS pulse intensity (in A/μs) to activate a minimum surface of 4 mm^2^ of left lobule VIII (i.e., VIII-A and VIII-B combined) in each head model (i.e., Atlas 1-5) for different coil types and locations.

| Coil Type | Location | TMS pulse intensity (*A/μs*) to activate 4 mm<sup>2</sup> surface in left lobule VIII |  |  |  |  |
| --- | --- | --- | --- | --- | --- | --- |
|  |  | Atlas 1 | Atlas 2 | Atlas 3 | Atlas 4 | Atlas 5 |
| MagStim D70 | *3LII* | 285.94 | 248.86 | 479.45 | 318.78 | 257.42 |
|  | *L8p* | 272.56 | 232.77 | 215.50 | 250.02 | 184.39 |
| Deymed 120BFV | *3LII* | 201.37 | 161.44 | 201.75 | 181.18 | 164.41 |
|  | *L8p* | 185.39 | 132.67 | 165.12 | 164.55 | 146.78 |

For TMS pulses at intensities in **Table 2-3**, the predicted activation site of Purkinje cells was primarily the axon *proximal* segment (>99% of activated cells in 15 out of 20 values in **Table 2** and 14 out of 20 values in **Table 3**; min: 69.53% and 65.14%, respectively), while the other segments were rarely activated as first (i.e., soma: >10% of activated cells in 3 out of 20 values and 4 out of 20 values, respectively; axon *intermediate* segment: >10% of activated cells in 2 out of 20 values and 1 out of 20 values, respectively; axon *terminal* segment: <0.001% of activated cells across all values in **Table 2-3**). More importantly, the spillover effects to non-motor-related lobules, i.e., left Crus I-II and VII-B, significantly decreased for coils positioned in *L8p* compared to *3L1I* when the pulse intensity is kept at the values in **Table 2-3**. Specifically, for each configuration and pulse intensity in the tables, we calculated the ratio between the activated surface area in lobule VII (i.e., combination of Crus I-II and VII-B) and lobule VIII, and we found that such ratio decreased by 74.05±17.26% and 78.53±18.66% for MagStim D70 and Deymed 120BFV coils, respectively, for intensities in **Table 2**, and 70.61±16.27% and 71.74±20.83%, respectively, for intensities in **Table 3**. No significant spillover (i.e., activated surface area less than 0.01 mm^2^) to the left lobule V-VI or the right hemisphere, instead, was reported for any configuration.

Since these results were obtained for currents flowing downwards in the coil, we also reversed the coil current direction. We obtained a mild but consistent reduction in the activation of lobule VIII and increased activation of lobule VII, which were reflected in higher TMS intensities compared to the values in **Table 2-3**, see ***Supplementary Information*, Table S5-S6**, respectively. However, we did observe a consistent shift in the distribution of activated Purkinje cells due to the change in E-field polarity.

Altogether, these results indicate that the optimized positions *L8p* can result in a stronger neuronal response in the target ROI while significantly decreasing the spillover effects to the neighboring structures. When the left lobule VIII is the target ROI, this can lead to more selective activation of the cerebrocerebellar area with modest secondary recruitment of spinocerebellar fibers.

## DISCUSSION

Cerebellar TMS is commonly used to investigate cerebello-cerebral interactions and has been recently proposed to treat cerebellar disorders. The effect size of cerebellar TMS, though, is regarded as low [12], and inconsistent outcomes have been frequently reported, especially when the coil is at the *3L1I* fiduciary marker [7, 65, 66]. The scalp-to-cerebellum distance, which is approximately twice the scalp-to-cerebrum distance [9], is thought to strongly reduce the neuronal response to magnetic pulses compared to cerebral TMS [62]. Studies [67–69], though, have shown that the functional response to cerebellar TMS can dramatically increase when the coil is moved away from the *3L1I* position and towards the parietal lobe, which suggests that alternative coil placements can significantly increase the effect size of TMS. However, there is a lack of rationale for these positions, partly because the spatial extent of the response of cerebellar neurons and fibers to TMS-induced E-fields remains unclear.

Our study provides new computational tools to rapidly explore alternative coil positions and types (i.e., nonplanar double-cone) and correspondingly predict the magnitude and spatial arrangement of the cellular responses to TMS pulses. Although limited to Purkinje cells for now, our solution accounts for the cellular morphology, which critically shapes the response of pyramidal cells to cerebral TMS, e.g., [48, 64, 70], and provides a realistic representation of the neural bundles within the white matter, including the nonlinear shape and orientation of the axonal fibers. Also, while the coil placement optimization methods for cerebral TMS assume tangency to a convex scalp surface at the target ROI, e.g., [17–19, 24–26], our solution fully accounts for the nonconvex geometry of the human posterior neck region and precisely calculates the arrangement of the E-fields within the cerebellar volume. Finally, our activation threshold prediction algorithm based on GPR models requires the estimation of just a few model parameters, which makes our tool easy to train on small datasets. Perhaps more importantly, our results demonstrate high robustness against morphological variability across subjects and consistently low prediction errors on new data. This is a significant departure from prior solutions for model-based activation threshold estimation, e.g., [33, 34], which primarily focused on simplified axonal geometries and current source models. More recently, artificial neural networks have been shown to estimate the activation threshold of neurons based on E-field measurements with low prediction errors [35, 36]. These algorithms, however, require one network model per neuron template, and multiple neuron templates are needed to recapitulate different cellular morphological properties, which make these algorithms hard to train and difficult to scale-up as the number of cell morphologies increases. Our solution, instead, is independent of cell morphologies, can be extended to any cell model with similar ion channel composition, and is trained quickly.

When applied to cerebellar TMS, our approach showed that, albeit the spatial component of the E-field is inversely correlated with the Purkinje cell activation threshold (***Supplementary Information*, Fig. S4-5**), the magnitude of the E-field at the cerebellar surface is hardly indicative of action potentials in efferent fibers. Moreover, although the E-field distribution for any TMS setup varied modestly across individuals, which is consistent with prior work [16], we found that the amount of cellular activation can change significantly across subjects as a result of inter-subject differences in lobule size and adjustments to the coil orientation. Also, we found that the activation site on Purkinje cells can vary from one lobule to the next and generally moves towards the axon distal segments as the distance from the coil increases. In our models, in fact, the soma and axon *proximal* segment were the primary activation sites in outer lobules, e.g., VII-B and Crus I-II, for both coils, whereas the axon *terminal* segment was the primary activation site in lobules VIII-B (both coil types) and V-VI (Deymed 120BFV coil only), albeit high pulse intensities were necessary to directly elicit an action potential in deeper regions. Finally, our calculations indicate that a coil position, i.e., *L8p*, further left of the position indicated in [67–69] can activate significantly more efferent fibers within lobule VIII-A and the entire cerebellar motor area while reducing the spillover effects both in adjacent areas (e.g., VII-B and Crus I-II) and the occipital lobe, which is expected to reduce nonmotor side effects of the stimulation. The effects on spinal mechanisms and peripheral nerves [71], though, were not explicitly accounted for in our study and may require experimental validation.

Although we built our algorithms on geometrically realistic models of Purkinje cells and MRI-derived head models, we acknowledge that several limitations may apply to the computational pipeline. First, we note that the activation thresholds were measured by temporarily disabling the Purkinje cells’ spontaneous firing despite these neurons being tonically active at rest. This may have affected the spatial arrangement of the estimated activation thresholds because the firing rate at rest is thought to modulate the response of Purkinje cells to exogenous stimuli [72, 73]. The net effect across the entire cerebellum volume, though, is likely modest due to the limited extent of the coordination between the Purkinje cell firing across lobules.

Secondly, we note that the limited resolution of the 3T MRI images led to a lack of segmentation for the dentate, interposed, and fastigial nuclei in the corpus medullare and coarser grey matter-white matter boundaries in the vermis region, which might have negatively affected the shape of some axon fibers. Although this may be addressed in the future by including cerebellar tractography [20–22, 74], the impact of the MRI resolution on the estimated activation thresholds is expected to be modest because we limited our reconstruction of the Purkinje cell projections to the primary axon. While axon collaterals are known to play an important role in cell-to-cell connectivity within the Purkinje layer [30], Purkinje cell primary axons form the bulk of the fiber bundles towards the cerebellar nuclei [59]. Moreover, the axonal collaterals are unmyelinated [75], which makes them hard to activate with the tested TMS settings, whereas the polarization at the axon bend is known to facilitate the initiation of action potentials [76].

Finally, our Purkinje cell models include soma, dendrites, and axon stem from [54], which were derived from a guinea pig preparation and are smaller in size compared to the human Purkinje cells [77, 78]. The size differences suggest that our predictions might overestimate the actual activation thresholds, as larger neurons tend to be more excitable by extracellular E-fields [79]. A recent study with multicompartmental pyramidal neuron models [64], though, indicate the activation threshold tends to decrease proportionally to the increment of the neuron’s size, and therefore activation thresholds for human Purkinje cells could be extrapolated from our predictions. Perhaps more interestingly, since the Purkinje cell somas and axon stems are approximately twice the size in humans compared to guinea pigs [77, 78], the extrapolated activation thresholds would indicate that cerebellar TMS can evoke wide neural activation for pulse intensities well below the maximum stimulator outputs, i.e., 100-120 A/µs, calculated in [39] for both coils.

## CONCLUSIONS

Our study developed a computational pipeline that optimizes the coil placement and predict the neuronal responses to TMS-induced E-fields in the cerebellum. These tools facilitate further investigation of the neural mechanisms underlying cerebellar TMS and provide rationally designed coil placements to help improve the outcomes of future cerebellar TMS studies.

## ACKNOWLEDGEMENT

This work was partly supported by the US NSF (National Science Foundation) CAREER Award 1845348 to S.S. The funders had no role in study design, data collection and analysis, decision to publish, or preparation of the manuscript. The authors thank Jianghua Wu at CUHK-Shenzhen and Fan Jiang at the Georgia Institute of Technology for helpful discussions on machine learning and optimization procedures.

## DATA AVAILABILITY STATEMENT

The code and relevant data of this study will be made available on GitHub upon acceptance.

