## Supplementary Information for "Computational Methods for Optimal Coil Placement and Maximization of Lobule-Focused Cortical Activation in Cerebellar TMS"

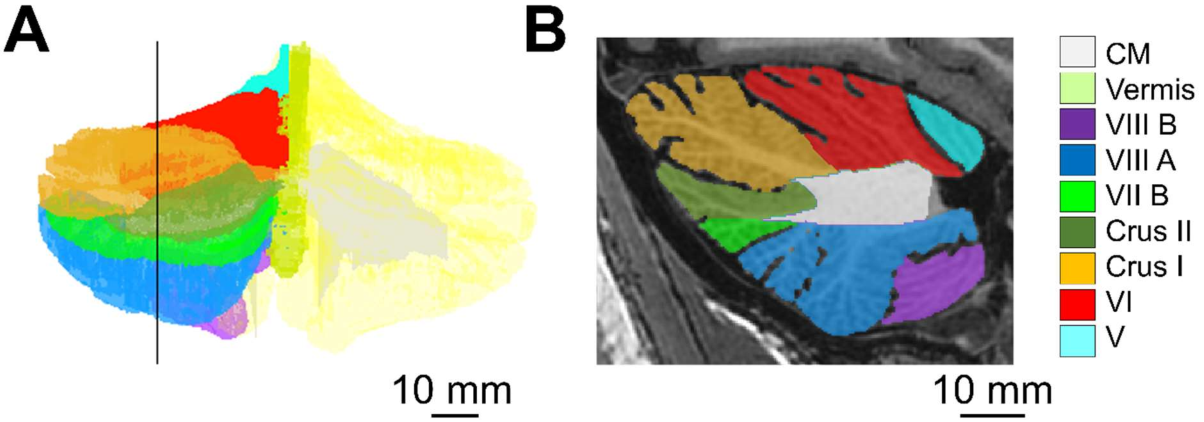


**Figure S1. A-B)** 3D view (**A**) and sagittal view (**B**) of the segmented cerebellar lobules from one head model in [1] (Atlas 5). The slice depicted in **B)** corresponds to the vertical black line in **A)**. The color legend on the right denotes the position of the corpus medullare (CM) and lobules V-VIII, including the sub-lobule regions Crus I, Crus II, VII-B, VIII-A, and VIII-B, in both panels. Panel **A)** also depicts the position of the vermis (*light green*), which resulted from the application of the ACAPULCO segmentation software [2].

| Tissue | Mean Conductivity (*S/m*) |
|---|---|
| White Matter | 0.126 |
| Gray Matter | 0.275 |
| Cerebrospinal Fluid | 1.654 |
| Scalp | 0.465 |
| Eyes | 0.5 |
| Compact Bone | 0.008 |
| Spongy Bone | 0.025 |
| Blood | 0.6 |
| Muscle | 0.16 |

**Table S1.** Mean conductivity value for the types of tissue segmented in the head models.

| Atlas | Coil type | Position | Coil center (*mm*) | | | Coil orientation (*mm*) | | |
|---|---|---|---|---|---|---|---|---|
| | | | *x* | *y* | *z* | *x* | *y* | *z* |
| 1 | MagStim D70 | *3L1I* | -29.20 | -110.19 | -40.48 | 0.07 | -0.31 | 0.95 |
| | | *L8p* | -64.05 | -78.36 | -51.94 | -0.54 | 0.40 | 0.74 |
| | Deymed 120 BFV | *3L1I* | -38.42 | -131.84 | -43.90 | -0.04 | 0.16 | -0.99 |
| | | *L8p* | -72.73 | -105.63 | -56.93 | 0.14 | 0.06 | -0.99 |
| 2 | MagStim D70 | *3L1I* | -29.54 | -108.45 | -38.34 | 0.02 | -0.17 | 0.99 |
| | | *L8p* | -65.42 | -75.20 | -53.35 | -0.46 | 0.58 | 0.67 |
| | Deymed 120 BFV | *3L1I* | -34.57 | -124.55 | -41.28 | -0.03 | 0.12 | -0.99 |
| | | *L8p* | -64.48 | -102.76 | -56.48 | 0.05 | 0.09 | -0.99 |
| 3 | MagStim D70 | *3L1I* | -24.77 | -110.91 | -39.70 | 0.53 | -0.42 | 0.73 |
| | | *L8p* | -55.00 | -78.20 | -52.81 | -0.02 | -0.45 | 0.89 |
| | Deymed 120 BFV | *3L1I* | -29.73 | -128.16 | -42.30 | -0.02 | 0.30 | -0.95 |
| | | *L8p* | -63.96 | -108.83 | -70.87 | 0.10 | 0.32 | -0.94 |
| 4 | MagStim D70 | *3L1I* | -26.99 | -108.39 | -38.57 | 0.29 | -0.35 | 0.89 |
| | | *L8p* | -66.18 | -76.08 | -51.69 | -0.31 | -0.07 | 0.95 |
| | Deymed 120 BFV | *3L1I* | -34.56 | -129.79 | -42.89 | 0.06 | 0.18 | -0.98 |
| | | *L8p* | -76.92 | -97.17 | -53.00 | 0.15 | 0.17 | -0.97 |
| 5 | MagStim D70 | *3L1I* | -29.12 | -109.32 | -42.58 | 0.34 | -0.38 | 0.86 |
| | | *L8p* | -67.82 | -69.82 | -58.68 | -0.38 | 0.18 | 0.91 |
| | Deymed 120 BFV | *3L1I* | -38.69 | -125.46 | -46.94 | 0.14 | 0.10 | -0.99 |
| | | *L8p* | -78.48 | -94.92 | -46.45 | 0.08 | 0.12 | -0.99 |

**Table S2.** Coordinates (*x*, *y*, *z*) of the coil's center point (*Coil center*) and orientation (*Coil orientation*) for two coil configurations, i.e., *3L1I* and *L8p* (see definition in the *Main Text*), calculated for each head model in [1], i.e., Atlas 1, 2, 3, 4, and 5. Configurations *3L1I* and *L8p* were computed for MagStim D70 and Deymed 120BFV coils and are reported in the MNI coordinate system. The MNI coordinate system is depicted in **Fig. 1C** in the *Main Text* and refers to the MNI 152 template [3]. The coil's orientation is expressed by reporting the projections of the coil's transverse unit vector $\hat{\mathfrak{t}}$ (*green arrows* in **Fig. 1C** in the *Main Text*) along the axes of the MNI system, with $\hat{\mathfrak{t}}$ placed at the coil's center point. Note that MagStim D70 and Deymed 120BFV coils are oriented roughly in opposite directions in each configuration because of the opposite directions of the current flow.

### 1. *Training and Testing of the Fast Purkinje Cell Activation Threshold Prediction Algorithm*

A total of 200,000 Purkinje cell (PC) models (40,000 per head model) were used to cross-validate the algorithm for fast Purkinje cell activation threshold prediction. The activation thresholds (in A/µs) of these PC models were calculated as described in the *Main Text* via a binary search algorithm with 0.5 A/µs accuracy when a Deymed 120BFV coil is adopted and placed in positions *L8p*, see **Table S2**. In each head model, the locations of the 40,000 Purkinje cells were uniformly distributed across the left lobules V (5,000 locations), VI (5,000 locations), VII (15,000 locations), and VIII (15,000 locations). For each PC model, it was also annotated whether the action potential started at the soma, *proximal* axon segment, *intermediate* axon segment, or *terminal* axon segment when the activation threshold was met.

A five-fold cross-validation scheme was adopted, i.e., at each fold, the prediction algorithm was trained on the PC activation thresholds from one head model (*training set*) and tested on the activation thresholds of PC models from the remaining four head models (*test set*). The training phase consisted of estimating the parameters of the Gaussian process regression models and piecewise linear models (i.e., $GPR_w$, $w = S, P, I$ and $PWL_T$ in **Fig. 1B** in the *Main Text*) and was conducted separately for each model on the activation thresholds of those Purkinje cells whose action potential started at the soma (for $GPR_S$), *proximal* axon segment (for $GPR_P$), *intermediate* axon segment (for $GPR_I$), or *terminal* axon segment (for $PWL_T$), respectively, when the activation threshold was met.

To cope with the reduced number of training samples per model and obtain a balanced probing of the range of activation thresholds, the training set for each model (i.e., either GPR or PWL) was augmented via scaling. Briefly, for each model, the training pairs $(\mathbf{x}, y)$, where $\mathbf{x}$ is the feature vector and $y$ is the model's output (see **Fig. 1B** in the *Main Text*), were multiplied by random numbers uniformly distributed between 0.1 and 2, and up to 30,000 of the resultant scaled pairs were added to the training set. Augmentation through scaling was chosen because an inverse relationship was noted between the intensity of the *normal* E-fields along the PC compartments across the cerebellar mesh grid and the cell activation thresholds. Furthermore, the number of scaled pairs added to each training set (i.e., 20,000 for $GPR_S$ and $GPR_P$; 30,000 for $GPR_I$ and $PWL_T$) was determined to minimize the mean absolute prediction error on test data, and the selection of the scaled pairs was done to ensure that the resultant augmented training sets provide a balanced sample of the activation threshold values up to 600 A/µs.

The GPR model parameters were estimated in MATLAB, rel. 2021b, by using a quasi-Newton optimizer with trust-region method (function `fitgpr`, Statistics and Machine Learning Toolbox). The PWL model parameters, i.e., $k_1$ and $k_2$ in the *Main Text*, instead, were estimated as the median values of the ratio $\mathbf{x}/y$ for $\mathbf{x} \geq 0$ and $\mathbf{x} < 0$, respectively ($\mathbf{x}$ is a feature scalar for the *terminal* axon segment, see the *Main Text*).

The prediction performance of the individual models (i.e., GPR or PWL) and the final prediction rule

$\hat{A}_{thr} = \alpha/y_r$ (see **Fig. 1B** in the *Main Text*) were assessed on each test set in terms of mean absolute error (MAE) and mean absolute percentage error (MAPE). In addition, for each test set and neuron segment, a linear regressor was fit on the pairs formed by the simulated activation thresholds for the neuron segment (independent variable) and the thresholds provided by the correspondent prediction model (dependent values), and the coefficient of determination, $R^2$, was used to measure the goodness-of-fit.

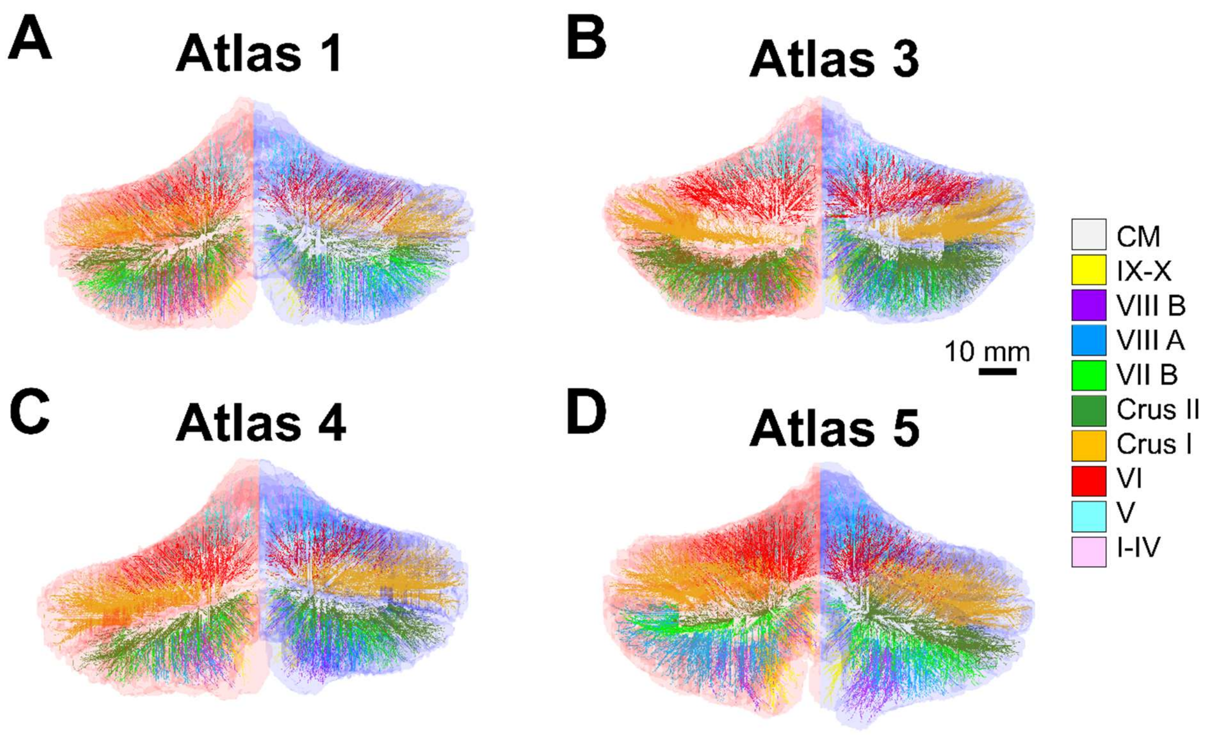


**Figure S2. A-D)** 3D geometry of the multicompartmental axons attached to the Purkinje cell models in four distinct head models from [1], i.e., Atlas 1 (**A**), Atlas 3 (**B**), Atlas 4 (**C**), and Atlas 5 (**D**). Note that only 1% of the estimated axons per head model is depicted. Axons in all panels are colored according to the legend on the right side to indicate the lobules from which they depart. CM: corpus medullare; I-X: lobules I-X, including the sub-lobule regions Crus I, Crus II, VII-B, VIII-A, and VIII-B.

| Prediction Model | Test Set 1 | Test Set 2 | Test Set 3 | Test Set 4 | Test Set 5 |
|---|---|---|---|---|---|
| $GPR_S$ | 0.986 | 0.994 | 0.989 | 0.996 | 0.995 |
| $GPR_P$ | 0.994 | 0.996 | 0.995 | 0.998 | 0.997 |
| $GPR_I$ | 0.985 | 0.985 | 0.985 | 0.985 | 0.985 |
| $PWL_T$ | 0.991 | 0.993 | 0.991 | 0.992 | 0.995 |
| $\hat{A}_{thr}$ | 0.989 | 0.994 | 0.992 | 0.995 | 0.994 |

**Table S3.** Goodness-of-fit of the GPR models, PWL models, and prediction rule $\hat{A}_{thr}$ on simulated activation thresholds. The coefficient of determination ($R^2$) of the linear regressor between the simulated activation threshold and the predicted activation threshold on test data (*Test Set*) is reported. Test Sets 1, 2, 3, 4, and 5 include data from head models not used for training. For each Test Set, the training was conducted on simulated Purkinje cell activation thresholds from Atlas 1, 2, 3, 4, and 5 in [1], respectively.

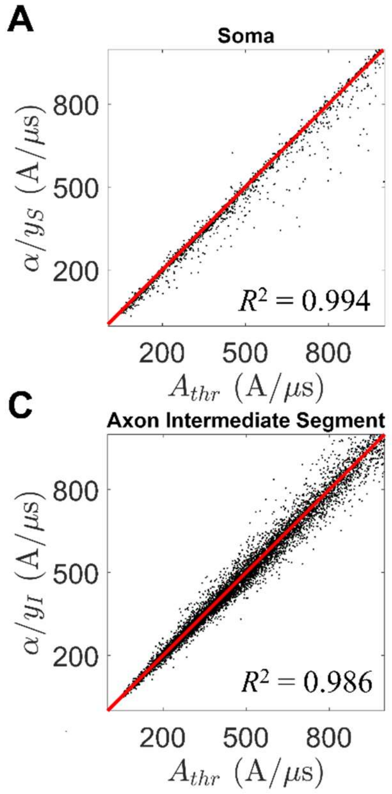


**Figure S3. A-D)** Activation threshold $\alpha/y_r$ predicted by the GPR model ($r = S, P, I$) or the PWL model ($r = T$) versus the actual activation threshold $A_{thr}$. Symbols $\alpha$ and $y_r$ ($r = S, P, I, T$) are like in **Fig. 1B** in the *Main Text*. In each panel, black dots denote PC models from Test Set 2 for which the TMS-induced action potentials start at the soma (**A**), axon *proximal* segment (**B**), axon *intermediate* segment (**C**), or axon *terminal* segment (**D**), respectively, and therefore their activation thresholds are predicted by the model $GPR_S$, $GPR_P$, $GPR_I$, or $PWL_T$, respectively. In each panel, a linear regressor (*red* line) is fit on the black dots, and the goodness-of-fit is provided by the coefficient of determination, $R^2$.

| Prediction Model | Test Set 1 | Test Set 2 | Test Set 3 | Test Set 4 | Test Set 5 |
|---|---|---|---|---|---|
| $GPR_S$ | 4.188 (1.517%) | 2.621 (1.002%) | 4.141 (1.538%) | 2.441 (0.924%) | 2.700 (1.069%) |
| $GPR_P$ | 3.579 (1.316%) | 2.566 (0.903%) | 3.245 (1.198%) | 2.617 (0.966%) | 2.859 (0.983%) |
| $GPR_I$ | 12.984 (4.113%) | 12.817 (3.906%) | 12.939 (4.076%) | 12.626 (3.939%) | 12.870 (3.936%) |
| $PWL_T$ | 4.591 (1.504%) | 4.194 (1.289%) | 4.485 (1.491%) | 4.440 (1.408%) | 4.186 (1.329%) |
| $\hat{A}_{thr}$ | 5.585 (1.861%) | 4.419 (1.395%) | 5.187 (1.773%) | 4.462 (1.457%) | 4.890 (1.587%) |

**Table S4.** Five-fold cross-validation of the GPR models, PWL models, and prediction rule $\hat{A}_{thr}$. For each fold, the mean absolute error (MAE, in A/µs) and mean absolute percentage error (MAPE, in %) are calculated on test data (*Test Set*). Test Sets 1, 2, 3, 4, and 5 include data from head models not used for training. For each Test Set, the training was conducted on simulated Purkinje cell activation thresholds from Atlas 1, 2, 3, 4, and 5 in [1], respectively. Values are reported as MAE (MAPE).

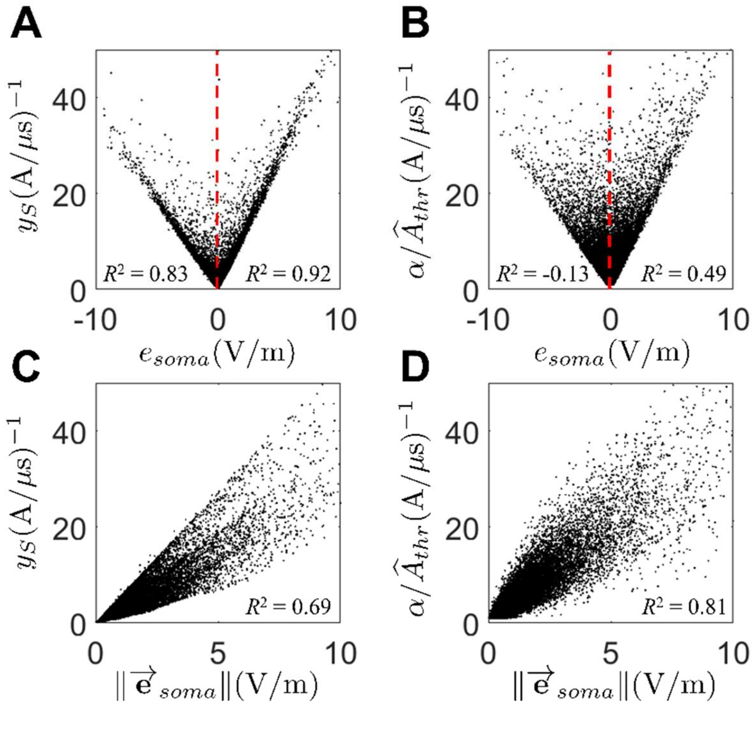


**Figure S4. A-B)** Output, $y_S$, of the peri-somatic GPR model (i.e., $GPR_S$ in the *Main Text*) (**A**) and scaled reciprocal of the predicted activation threshold, $\hat{A}_{thr}$ (**B**) versus the projection, $e_{soma}$, of the E-field along the somato-axonal axis of the Purkinje cells. **C-D)** Outputs $y_S$ (**C**) and $\alpha/\hat{A}_{thr}$ (**D**) versus the magnitude, $\|\vec{e}_{soma}\|$, of the E-field at the somatic compartment. In **B)** and **D)**, $\alpha = 10^3$ is like in **Fig. 1B** in the *Main Text*. In each panel, linear regressors are fit on the positive black dots and the negative black dots (if any), and the goodness-of-fit is provided by the coefficient of determination, $R^2$, on the right side and left side, respectively. In each panel, data points from Test Set 2 are depicted.

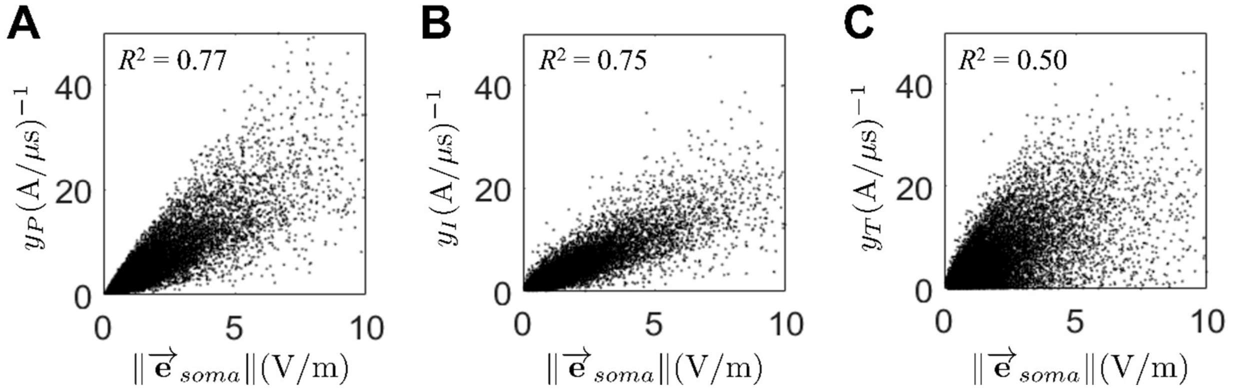


**Figure S5. A-C)** Output, $y_P$, of the axon *proximal* GPR model (i.e., $GPR_P$ in the *Main Text*) (**A**), output, $y_I$, of the axon *intermediate* GPR model (i.e., $GPR_I$ in the *Main Text*) (**B**), and output, , $y_T$, of the axon *terminal* PWL model (i.e., $PWL_T$ in the *Main Text*) (**C**) versus the magnitude, $\|\vec{e}_{soma}\|$, of the E-field at the somatic compartment. In each panel, a linear regressor is fit on the black dots, and the goodness-of-fit is provided by the coefficient of determination, $R^2$, on the top left side. In each panel, data points from Test Set 2 are depicted.

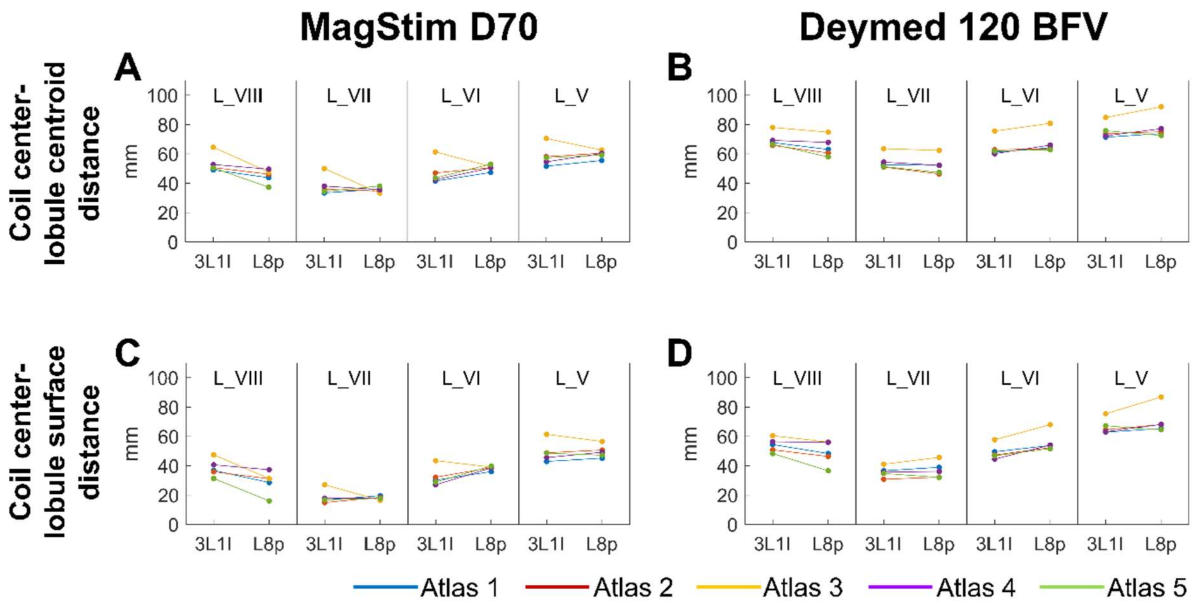


**Figure S6**. **A-B)** Euclidean distance between the coil's center point and the centroid of the left lobule V (L_V), VI (L_VI), VII (L_VII, including VII-B, Crus I, and Crus II), and VIII (L_VIII, including VIII-A and VIII-B) in each head model from [1] when the coil is the MagStim D70 (**A**) or the Deymed 120 BFV (**B**) in position *3L1I* or *L8p*. The average distance of the coil's center point from the centroid of L_VIII (i.e., target ROI) is 44.93±4.74 mm (MagStim D70) and 64.87±6.66 mm (Deymed 120BFV) when the coil is in position *L8p*, and 53.51±6.30 mm (MagStim D70) and 69.47±4.98 mm (Deymed 120BFV) when the coil is in position *3L1I* (mean±S.D. across five head models). **C-D)** Minimum distance between the coil's center point and the outer surface of lobules V-VIII in each head model when the coil is the MagStim D70 (**C**) or the Deymed 120BFV (**D**) in position *3L1I* or *L8p*. The minimum distance between the coil's center point and the outer surface of L_VIII is, on average, 28.86±7.85 mm (MagStim D70) and 48.69±8.07 mm (Deymed 120BFV) when the coil is in position *L8p*, and 38.43±6.04 mm (MagStim D70) and 54.10±4.76 mm (Deymed 120BFV) when the coil is in position *3L1I* (mean±S.D. across five head models). The color legend at the bottom of the figure applies to **A-D)**. Distances calculated for positions *L8p* and *3L1I* were statistically different (Wilcoxon rank sum test; *P*-value $P<0.01$) in case of L_VIII. No statistically significant difference was found between distances calculated for positions *L8p* and *3L1I* in case of L_V, L_VI, and L_VII (Wilcoxon rank sum test; $P>0.05$).

| Coil Type | Location | TMS pulse intensity (A/µs) to activate 1 $mm^2$ surface of left lobule VIII | | | | |
|---|---|---|---|---|---|---|
| | | **Atlas 1** | **Atlas 2** | **Atlas 3** | **Atlas 4** | **Atlas 5** |
| MagStim D70 | *3L1I* | 320.06 | 272.86 | 474.62 | 347.06 | 223.61 |
| | *L8p* | 277.54 | 264.97 | 271.27 | 277.59 | 146.00 |
| Deymed 120BFV | *3L1I* | 234.60 | 173.69 | 253.61 | 204.93 | 163.01 |
| | *L8p* | 201.71 | 150.24 | 213.96 | 187.87 | 124.01 |

**Table S5.** TMS pulse intensity (in A/µs) to activate a minimum surface of 1 $mm^2$ of left lobule VIII (i.e., VIII-A and VIII-B combined) in each head model (Atlas 1-5) from [1] for different coil types and locations. Coil current direction is reversed compared to **Table 2** in the *Main Text*.

| Coil Type | Location | TMS pulse intensity (A/µs) to activate 4 $mm^2$ surface of left lobule VIII | | | | |
|---|---|---|---|---|---|---|
| | | **Atlas 1** | **Atlas 2** | **Atlas 3** | **Atlas 4** | **Atlas 5** |
| MagStim D70 | *3L1I* | 334.86 | 299.54 | 534.95 | 364.67 | 241.88 |
| | *L8p* | 300.49 | 302.02 | 289.51 | 297.46 | 140.49 |
| Deymed 120BFV | *3L1I* | 242.40 | 188.74 | 272.78 | 214.70 | 180.84 |
| | *L8p* | 215.99 | 161.25 | 228.20 | 200.45 | 135.58 |

**Table S6.** TMS pulse intensity (in A/µs) to activate a minimum surface of 4 $mm^2$ of left lobule VIII (i.e., VIII-A and VIII-B combined) in each head model (Atlas 1-5) from [1] for different coil types and locations. Coil current direction is reversed compared to **Table 3** in the *Main Text*.

### CITED REFERENCES


[1] Park MT, Pipitone J, Baer LH, Winterburn JL, Shah Y, Chavez S, et al. Derivation of high-resolution MRI atlases of the human cerebellum at 3T and segmentation using multiple automatically generated templates. Neuroimage 2014;95:217-31.

[2] Han S, Carass A, He Y, Prince JL. Automatic cerebellum anatomical parcellation using U-Net with locally constrained optimization. Neuroimage 2020;218:116819.

[3] Dadar M, Manera AL, Fonov VS, Ducharme S, Collins DL. MNI-FTD templates, unbiased average templates of frontotemporal dementia variants. Sci Data 2021;8(1):222.
